# CRTAC1-A reprograms extracellular matrix viscoelasticity to constrain glioma progression

**DOI:** 10.64898/2026.05.06.723198

**Authors:** Varinder Singh, Devansh Swadia, Pritam Saha, Debasish Nath, Nilesh Deokate, Sabyasachi Rakshit

**Affiliations:** Department of Chemical Science, Indian Institute of Science Education and Research Mohali, Punjab-140306; Institute of Nanoscience and Technology, Mohali, Punjab-140306

**Keywords:** CRTAC1-A, Tumor Microenvironment, Tumor mechanobiology, Dynamic Matrix Mechanics, Gliomas

## Abstract

Mechanical remodeling of the extracellular matrix (ECM) influences glioma progression, yet the molecular regulators that control tumor matrix mechanics remain poorly understood. By comparing low-grade gliomas (LGGs), associated with improved patient survival, with glioblastomas (GBMs), which carry a poor prognosis, we identified the cartilage-derived ECM protein CRTAC1-A as enriched in LGGs and significantly reduced in GBMs, with elevated expression correlating with improved patient survival. Restoration of CRTAC1-A suppressed glioma cell proliferation and invasion and enhanced temozolomide efficacy in three-dimensional tumor spheroid models. Mechanistically, CRTAC1-A directly interacts with collagen I and reorganizes collagen networks across multiple length scales, generating a mechanically compliant yet structurally resilient ECM with reduced stiffness, enhanced elastic recovery, and resistance to persistent remodeling. This viscoelastic normalization limits invasive remodeling while preserving matrix permeability and drug penetration. Together, these findings identify CRTAC1-A as a reversible regulator of tumor ECM mechanics that suppresses glioma malignancy.

## Introduction

Dynamic physical interactions between cancer cells and the ECM play a central role in tumor progression by shaping the mechanical landscape of the tumor microenvironment^1,2^. Gliomas, tumors of the central nervous system, account for a relatively small fraction of total cancers yet exert a disproportionate impact on survival and quality of life^3,4,5^. Among these, GBM (Grade IV) remains particularly lethal, with median survival measured in months despite intensive multimodal therapy^6^. Increasing evidence indicates that ECM remodeling intensifies with tumor grade, generating mechanically permissive microenvironments that promote invasive migration and therapeutic resistance^7^. In contrast, LGGs (Grades I–II) are comparatively slow-growing and associated with improved survival outcomes^8^. While progressive ECM stiffening and structural reorganization have been linked to malignant progression^9^, the molecular regulators that maintain tumor-restraining mechanical states in earlier disease stages remain poorly defined.

Addressing this question requires an integrative framework that combines molecular profiling with quantitative measurements of tissue mechanics. We therefore adopted an unbiased transcriptomic approach to identify genes exhibiting inverse expression patterns across glioma grades and prioritized candidates based on prognostic relevance^10,11^. This analysis identified Cartilage Acidic Protein 1 (*CRTAC1*) as significantly associated with improved survival in glioma patients. Notably, *CRTAC1* has previously been reported to suppress tumor progression in lung adenocarcinoma by modulating integrin–ECM interactions^12^. Given the established link between glioma aggressiveness and ECM mechanics, this association prompted us to investigate whether *CRTAC1* may function as a regulator of tumor matrix architecture.

*CRTAC1* encodes two major isoforms: CRTAC1-A, an extracellular protein, and CRTAC1-B, a membrane-associated isoform predominantly expressed in brain tissue. Isoform-specific analysis revealed that CRTAC1-A, rather than CRTAC1-B, is enriched in LGGs. This observation suggested that the extracellular isoform may contribute to the mechanically restrained phenotype of early-stage gliomas. We therefore hypothesized that CRTAC1-A functions as a reversible regulator of ECM organization, modulating viscoelastic properties that influence tumor evolution.

Here, we demonstrate that CRTAC1-A reorganizes collagen networks to generate a mechanically compliant yet structurally resilient ECM that limits invasive remodeling. By integrating transcriptomic analysis with multiscale mechanical characterization, we identify CRTAC1-A as a matricellular regulator that reprograms tumor ECM viscoelasticity, providing mechanistic insight into how molecular control of matrix mechanics can constrain glioma progression and influence therapeutic responsiveness.

## Results

### *CRTAC1* is enriched in LGG and predicts favourable survival

To identify ECM–associated regulators that may contribute to glioma progression, we analysed transcriptomic profiles from LGG and GBM patient cohorts using the GEPIA2 webserver, focusing on genes with inverse expression patterns. The inverse expression pattern was estimated for LGG versus normal controls and GBM versus normal controls using the LIMMA method, with thresholds set at log_2_ fold change (log_2_FC > 1) and false discovery rate (q < 0.01) (Supplementary Fig 1). This analysis identified 3,982 upregulated and 1,774 downregulated genes in LGG relative to normal controls, and 5,224 upregulated and 2,439 downregulated genes in GBM. Comparative evaluation revealed 23 genes exhibiting inverse expression patterns between LGG and GBM. Of these, 11 genes were significantly upregulated in LGG but downregulated in GBM, whereas 12 genes displayed the opposite pattern (Fig. 1a).

**Fig 1:**
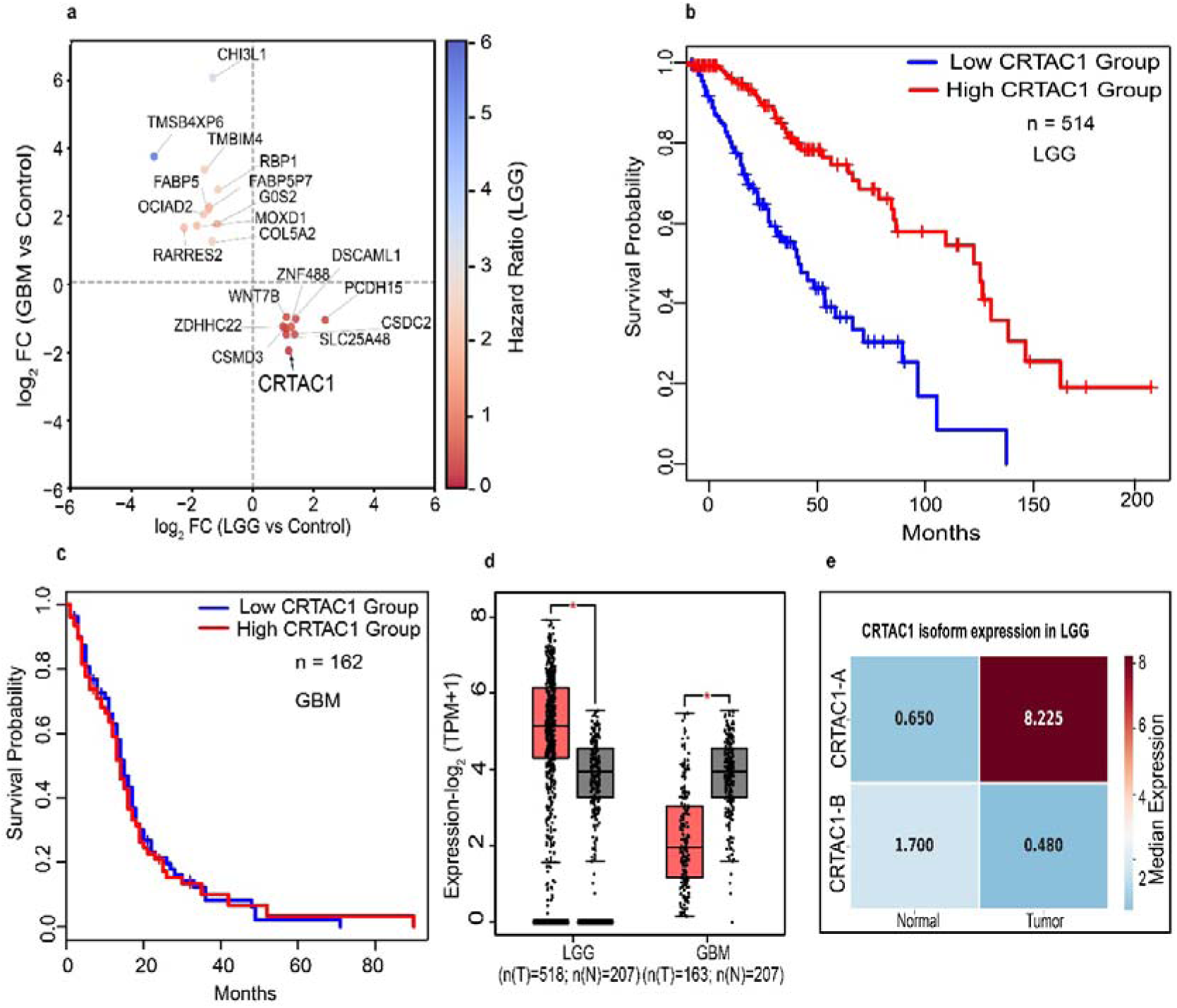
*CRTAC1* expression profiles and prognostic significance in LGG and GBM. (**a)** Differential expression analysis identifying 23 genes with inverse regulation patterns between LGG and GBM cohorts. Fold-changes were determined via the LIMMA method. Color-coding indicates the Hazard Ratio (HR), where blue denotes high-risk (increased HR) and red denotes low-risk (decreased HR) associations. (**b, c)** Kaplan-Meier survival curves stratified by *CRTAC1* expression in patients with LGG (**b**, n = 514) and GBM (**c**, n = 162). High *CRTAC1* expression is significantly associated with favorable outcomes in LGG (HR = 0.28; p < 0.0001), while no significant prognostic association is observed in GBM (HR = 1.0; p = 0.78). Statistical significance was assessed using the Log-rank (Mantel-Cox) test. (**d)** Comparative analysis of *CRTAC1* mRNA expression levels in LGG and GBM tumor tissues relative to non-malignant control subjects. (**e)** Median mRNA expression levels of CRTAC1-A and CRTAC1-B isoforms in LGG tumors, derived from the GEPIA2 database.

To determine prognostic relevance, Kaplan–Meier survival analyses were performed for all 23 candidates in both LGG and GBM cohorts (Supplementary Table 1). Among these, *CRTAC1* showed the strongest association with improved overall survival in LGG, with a hazard ratio for high expression of HR = 0.28 (p <0.0001) (Fig. 1b). In contrast, *CRTAC1* expression was markedly reduced in GBM (log□FC = −2.006; p <0.0001) (Fig. 1c). This inverse expression pattern across tumor grades, together with its strong prognostic association, identified *CRTAC1* as a candidate regulator of glioma progression (Fig. 1d).

*CRTAC1* encodes two major isoforms generated through alternative splicing: CRTAC1-A, an ECM-associated protein, and CRTAC1-B, a membrane-bound isoform predominantly expressed in brain tissue^13^. To determine which isoform accounts for elevated *CRTAC1* expression in LGG, we performed isoform-specific analysis using GEPIA2. CRTAC1-A was significantly upregulated in LGG compared with normal controls (log□FC = 2.483; p<0.0001) whereas CRTAC1-B was downregulated (log□FC = −0.867; p<0.0001) (Fig. 1e). These data indicate that increased *CRTAC1* expression in LGG is driven predominantly by the extracellular CRTAC1-A isoform rather than the brain-enriched CRTAC1-B isoform, suggesting a potential role for CRTAC1-A in the favorable clinical outcomes associated with LGG.

### Alternative splicing drives isoform switching from CRTAC1-B to CRTAC1-A in gliomas

To investigate the mechanism underlying CRTAC1-A upregulation in LGG, we analyzed percent spliced-in (PSI) values for the CRTAC1-A–specific exon using TCGA SpliceSeq data from LGG, GBM, and normal brain samples^14^ (Fig. 2a). PSI values reflect the relative inclusion of a given exon among all transcript isoforms.

**Fig 2:**
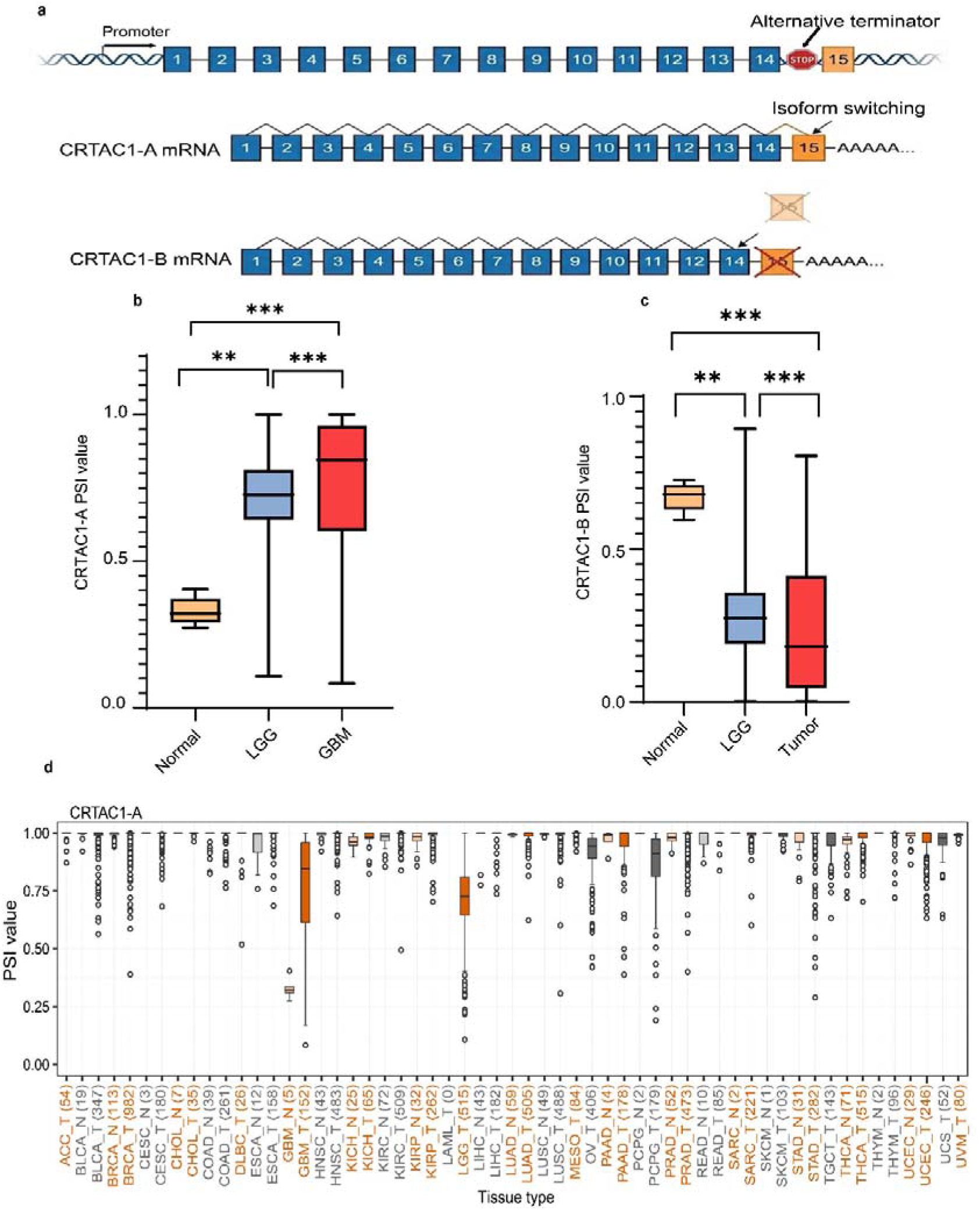
*CRTAC1* isoform switching and transcript-level dysregulation. (**a)** Schematic representation of alternative termination yielding the CRTAC1-A and CRTAC1-B isoforms. (**b, c)** Analysis of Percent Spliced-In (PSI) values for CRTAC1-A (**b**) and CRTAC1-B (**c**) in normal brain tissue (n = 5), LGG (n = 516), and GBM (n = 152). A reciprocal switching pattern is observed, characterized by significant CRTAC1-A enrichment and CRTAC1-B depletion in tumor samples relative to normal controls. (**d)** Pan-cancer analysis showing elevated CRTAC1-A PSI values across multiple tumor types using TCGA pan cancer database. For (**b)** and (**c)**, statistical significance was determined by two-tailed Mann-Whitney test (p < 0.001 for normal vs. LGG; p < 0.0001 for LGG vs. GBM and normal vs. GBM).

Both LGG and GBM tumors exhibited significantly higher PSI values for the CRTAC1-A specific exon compared to normal controls (median PSI: controls = 0.321; LGG = 0.726, p < 0.0001; GBM = 0.8426, p < 0.0002), indicating enhanced exon inclusion in gliomas. Moreover, PSI values were significantly higher in GBM than in LGG (Fig. 2b), suggesting progressive enrichment of CRTAC1-A favoring splicing with increasing tumor grade. In contrast, PSI values corresponding to CRTAC1-B–specific exons were reduced in both LGG and GBM relative to controls (Fig. 2c), consistent with a shift in isoform usage.

Importantly, although CRTAC1-A exon inclusion increases with tumor grade, overall CRTAC1-A transcript abundance declines in GBM due to progressive copy-number loss (Fig. 1d). These findings indicate that alternative splicing and transcript abundance are regulated through distinct mechanisms: exon inclusion favors CRTAC1-A in gliomas, whereas genomic deletion reduces total transcript levels in advanced disease. Analysis across multiple tumor types further revealed that enhanced CRTAC1-A exon inclusion is not restricted to gliomas but is observed broadly across cancers (Fig. 2d)^15^, suggesting a conserved splicing bias in malignant contexts.

### CRTAC1-A restrains glioma proliferation and invasion

Having identified CRTAC1-A as enriched in LGG and associated with favourable prognosis, we next examined its functional role in glioma cell behaviour. U87MG cells stably overexpressing CRTAC1-A (U87MG–CRTAC1A-His) were generated and compared with vector control cells (U87MG–pcDNA3.1). Proliferation was assessed using a 3-(4,5-dimethylthiazol-2-yl)-2,5-diphenyltetrazolium bromide (MTT assay over 120 h. We observed that CRTAC1-A overexpression reduced cell proliferation beginning at 96 h relative to controls (Fig. 3a), indicating that CRTAC1-A attenuates glioma cell growth under in vitro conditions.

**Fig 3:**
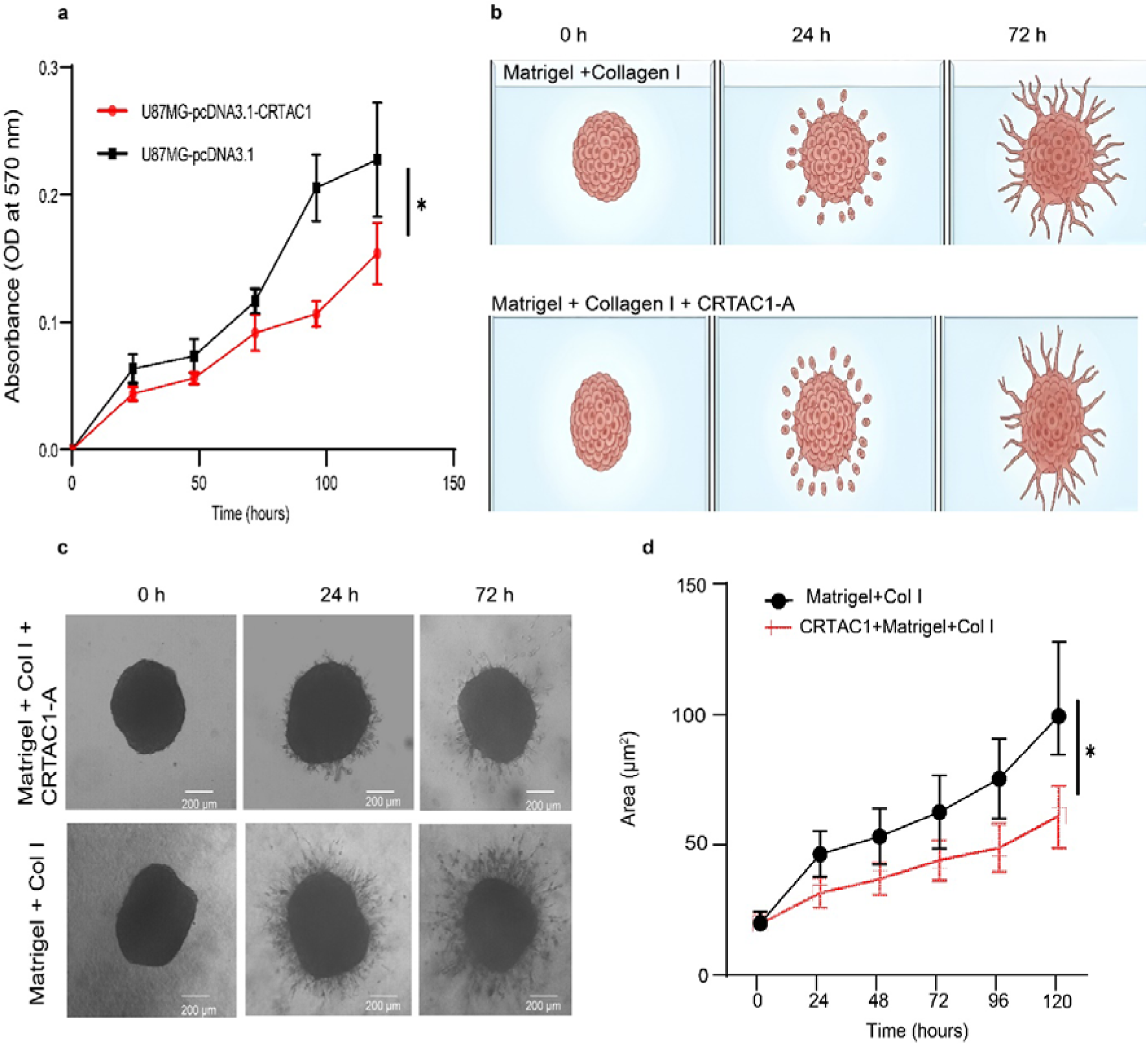
CRTAC1-A overexpression attenuates the proliferative and invasive capacity of U87MG cells. (**a)** Proliferation kinetics of U87MG cells with and without CRTAC1-A overexpression, monitored at 24-hour intervals over a 120-hour period (n = 6; p = 0.0143). (**b)** Schematic workflow of the 3D spheroid invasion assay. (**c)** Representative micrographs of U87MG spheroids embedded in Matrigel, captured at 0, 24, and 72 hours post-seeding in the presence or absence of CRTAC1-A (n = 23; 4x magnification; scale bar, 200 µm). (**d)** Quantitative analysis of invasive area, showing a significant reduction in invasive capacity following CRTAC1-A overexpression (n = 23; p= 0.0132). Statistical significance for (**a)** and (**d)** was determined by a two-tailed Mann-Whitney U-test.

Given that CRTAC1-A is an ECM-associated protein, we next investigated its impact on invasive behavior in a three-dimensional (3D) microenvironment. Recombinant CRTAC1-A was expressed in HEK293T cells, purified by affinity chromatography, and incorporated into reconstituted basement membrane (rBM; Matrigel). U87MG spheroids embedded in rBM alone exhibited robust radial invasion, whereas spheroids embedded in CRTAC1-A–supplemented matrices showed reduced invasive outgrowth (p = 0.0132) (Fig. 3b,c). The inhibitory effect of CRTAC1-A on invasion area became more pronounced over time; quantitative analysis showed a significant reduction of 35 µm^2^ at 96 hours, which further increased to a 50 µm^2^ difference by 120 hours relative to the control.

These findings indicate that extracellular CRTAC1-A limits glioma cell proliferation and suppresses invasive expansion in a 3D matrix context, supporting a tumor-restraining role linked to its ECM-modulatory function.

### IDH mutation and copy number alterations are associated with stage-dependent *CRTAC1* expression

Given the progressive reduction of *CRTAC1* expression across glioma grades, we next sought to identify genetic and epigenetic factors associated with its regulation. Mutations in Isocitrate Dehydrogenase (*IDH1* and *IDH2*) are characteristic of LGGs and occur less frequently in primary GBMs. Because *IDH* mutations are known to influence epigenetic states, we examined the relationship between *IDH* mutational status and CRTAC1-A expression in LGG tumors.

Compared with *IDH*-wildtype tumors, *CRTAC1* expression was significantly higher in *IDH1*-mutant (mean rank difference, MRD=191.5; p < 0.0001) and *IDH-2* mutant LGGs (MRD = 259.0; p < 0.0001), whereas IDH1 and IDH2 mutant tumors did not show a statistically significant difference among them (MRD = -67.40; p = 0.1246) (Fig. 4a). Consistent with this observation, ectopic expression of *IDH1*-R132H in U87MG cells increased *CRTAC1* transcript levels relative to *IDH1*-wildtype expression (log□FC = 6; p < 0.001) (Fig. 4b). These findings suggest that *IDH1* mutation status is associated with elevated *CRTAC1* expression in LGGs.

**Fig. 4:**
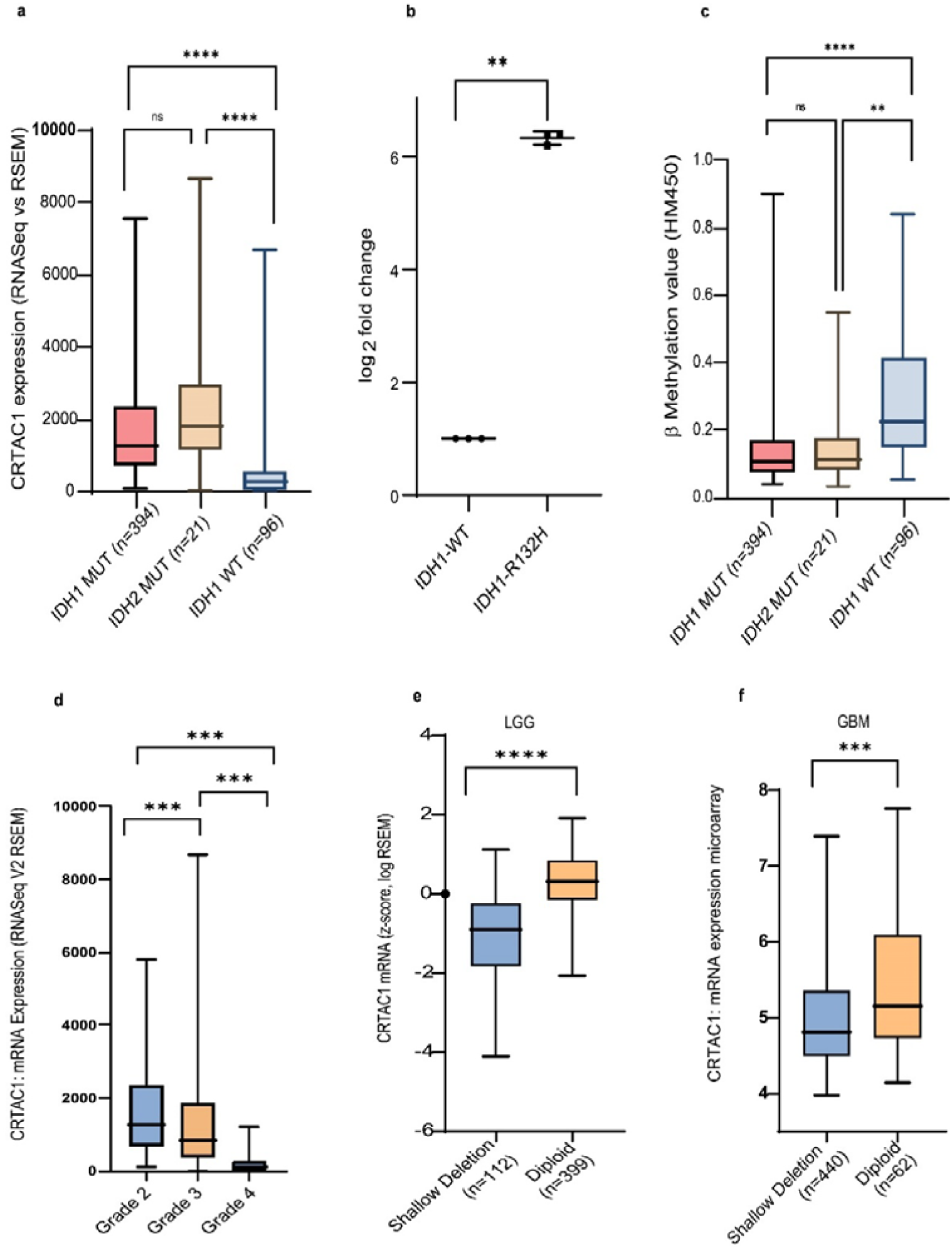
Epigenetic and genomic regulation of *CRTAC1* is associated with IDH1/2 mutational status and tumor grade. (**a)** Differential mRNA expression of *CRTAC1* across LGG cohorts stratified by IDH status. Significance determined by Kruskal-Wallis test: IDH1mt vs. IDH2mt (p = 0.1246); IDH1mt vs. IDHwt (p<0.0001); IDH2mt vs. IDHwt (p < 0.0001). (**b)** RT-qPCR analysis of *CRTAC1* mRNA levels in U87MG cells following ectopic expression of IDH1mt or IDH1wt constructs (n=3; p < 0.001). (**c)** DNA methylation β-values for the *CRTAC1* locus across IDH genotypes (IDH1mut vs. IDH2mut, p = 0.1746; IDH2mut vs. IDHwt, p < 0.001; IDH1mut vs. IDHwt, p < 0.0001). (**d)** Pan-glioma *CRTAC1* expression profiles across WHO tumor grades (Grades 2–4) sourced from cBioPortal (all pairwise comparisons, p < 0.0001). (**e, f)** Impact of *CRTAC1* shallow deletions on transcript abundance in LGG (**e**) and GBM (**f**) cohorts (p = 0.0002 for both). For **b-f**, statistical significance was determined using two-tailed Mann-Whitney test.

To determine whether *IDH* mutation also affects the splicing splicing switch favoring CRTAC1-A^16^, we compared CRTAC1-A PSI values between *IDH*-mutant and *IDH*-wildtype LGG samples. No significant difference was observed (Supplementary Fig. 2), indicating that alternative splicing toward the CRTAC1-A isoform occurs independently of IDH mutational status.

Because *IDH* mutations are linked to widespread epigenetic remodeling, we next examined *CRTAC1* promoter methylation. *IDH1* and *IDH2*-mutant LGG tumors exhibited reduced methylation at the *CRTAC1* promoter compared with *IDH*-wildtype tumors (MDF = □137.4; p < 0.0001, MDF = □127.4; p<0.001 respectively) (Fig. 4c), consistent with increased transcript abundance. Moreover, there was no significant methylation difference between *IDH1* and *IDH2*-mutant LGGs (MDF = □10.04; p>0.999). These data indicate that *IDH1*-associated epigenetic alterations may contribute to *CRTAC1* upregulation in LGGs.

Despite elevated expression in LGGs, *CRTAC1* levels declined progressively with increasing tumor grade (median grade 2= 1284, median grade 3= 814.1) and were lowest in GBM (median grade 4= 89.19) (p < 0.0001) (Fig. 4d). Given the high prevalence of copy number variations (CNVs) in gliomas, we assessed whether genomic alterations contribute to this reduction. CNV analysis from cBioPortal revealed that shallow deletions of the *CRTAC1* locus were associated with decreased transcript levels, with GBM tumors exhibiting the highest frequency of deletion and the lowest expression levels (p < 0.0001) (Fig. 4e, f).

Together, these findings suggest a stage-dependent regulatory model in which *IDH1* mutation–associated epigenetic modulation contributes to elevated *CRTAC1* expression in LGGs, whereas progressive copy-number loss is associated with reduced *CRTAC1* expression in GBMs. Thus, distinct genomic and epigenetic mechanisms appear to underlie the inverse expression pattern of *CRTAC1* across glioma grades.

### CRTAC1-A associates with collagen and reorganizes ECM architecture

CRTAC1-B has been reported to interact with the Nogo1 receptor, implicating it in neuronal plasticity regulation^17,18^. However, whether the extracellular isoform CRTAC1-A directly interacts with ECM components has remained unexplored. Given the central role of Collagen I in glioma-associated ECM, we first examined whether CRTAC1-A associates with collagen matrices.

Confocal imaging of GFP-tagged CRTAC1-A–secreting cells cultured on Cy5-labeled collagen demonstrated pronounced spatial colocalization between CRTAC1-A and collagen fibers (Fig. 5a, b). Quantitative analysis confirmed significant overlap, indicating incorporation or stable association of CRTAC1-A within the collagen network. To validate this interaction biochemically, we performed in vitro pull-down assays using purified recombinant proteins. Collagen I was retained in the presence of CRTAC1-A, while immunoblot analysis confirmed equivalent input across reactions (Supplementary Fig. 3). These findings establish a direct association between CRTAC1-A and collagen I.

**Figure 5:**
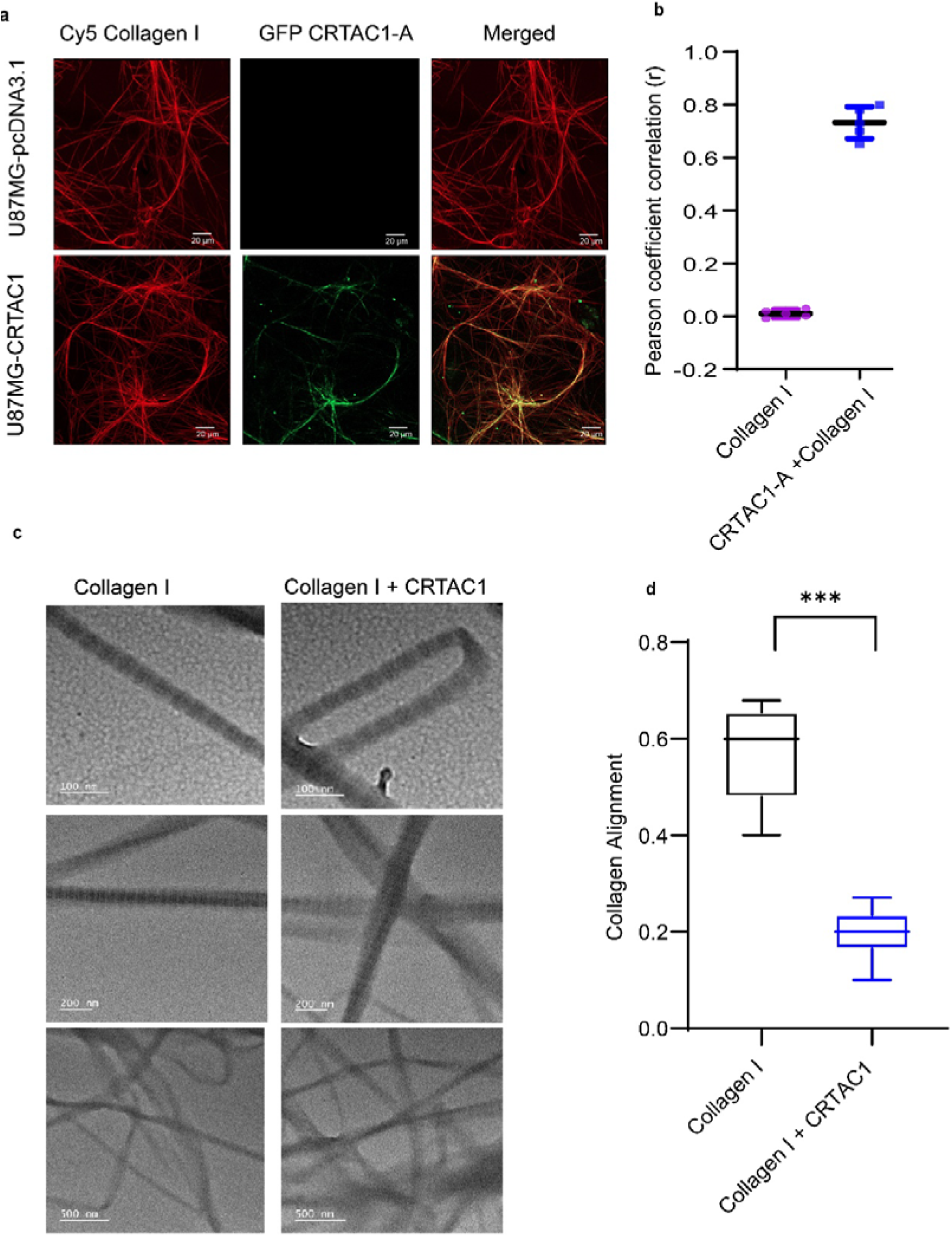
CRTAC1-A colocalizes with and modulates the structural organization of Collagen I. (**a)** Representative confocal immunofluorescence micrographs demonstrating the colocalization of CRTAC1-A and Collagen I. U87MG cells were transfected with either CRTAC1-A or empty pcDNA3.1 (control) and cultured on Cy5-labeled Collagen I matrices (n = 3; scale bar, 20 µm). (**b)** Quantification of colocalization using Pearson’s correlation coefficient. Values > 0.5 indicate a significant spatial correlation between CRTAC1-A and Collagen I. (**c)** Transmission electron microscopy (TEM) images of collagen fibrils polymerized in the presence or absence of CRTAC1 (n = 6). Scale bars represent 100 nm (top), 200 nm (middle), and 500 nm (bottom). (**d),** Quantitative assessment of fibril alignment and architecture. CRTAC1-A expression results in decreased linear alignment. Statistical significance was determined via two-tailed Mann-Whitney test.

To determine whether this interaction alters collagen organization, we next performed transmission electron microscopy (TEM) on reconstituted collagen I gels formed in the presence or absence of purified CRTAC1-A.

At low magnification, collagen-only matrices displayed a sparse fibrillar network with limited global connectivity and regions of low fibril density (Fig. 5c, lower panel). In contrast, matrices formed with CRTAC1-A exhibited increased fibrillar integration and enhanced network connectivity, indicating large-scale reorganization.

At intermediate magnification, collagen-only matrices were dominated by isolated fibrils with minimal sustained lateral association. In the presence of CRTAC1-A, fibrils formed laterally associated bundles extending over hundreds of nanometers and interconnected by junction-like nodes, reflecting enhanced interfibrillar connectivity (Fig. 5c, middle panel).

High-magnification imaging revealed that individual fibril ultrastructure and diameter remained preserved under both conditions, indicating that CRTAC1-A does not alter fibrillogenesis per se (Fig 5c, upper panel). Rather, quantitative analysis demonstrated reduced fibril alignment and increased junction density in CRTAC1-A–supplemented matrices (Fig. 5d), consistent with hierarchical reorganization of interfibrillar interactions.

Collectively, these multiscale ultrastructural analyses demonstrate that CRTAC1-A reorganizes collagen networks into a more interconnected and bundled architecture while preserving intrinsic fibril integrity. This structural remodeling provides a mechanistic basis for subsequent changes in ECM mechanical behavior.

### CRTAC1-A reprograms ECM mechanics

The hierarchical reorganization of collagen networks induced by CRTAC1-A prompted us to examine how these structural changes influence mesoscale matrix mechanics. To quantify bulk stiffness, we employed custom-built magnetic tweezers^19^ (Fig. 6a), in which embedded magnetic beads were subjected to controlled magnetic forces (Supplementary Fig. 4a,b). Matrix stiffness was determined from the force–displacement relationship by calculating the ratio of applied force to bead displacement (Fig. 6b).

**Fig 6:**
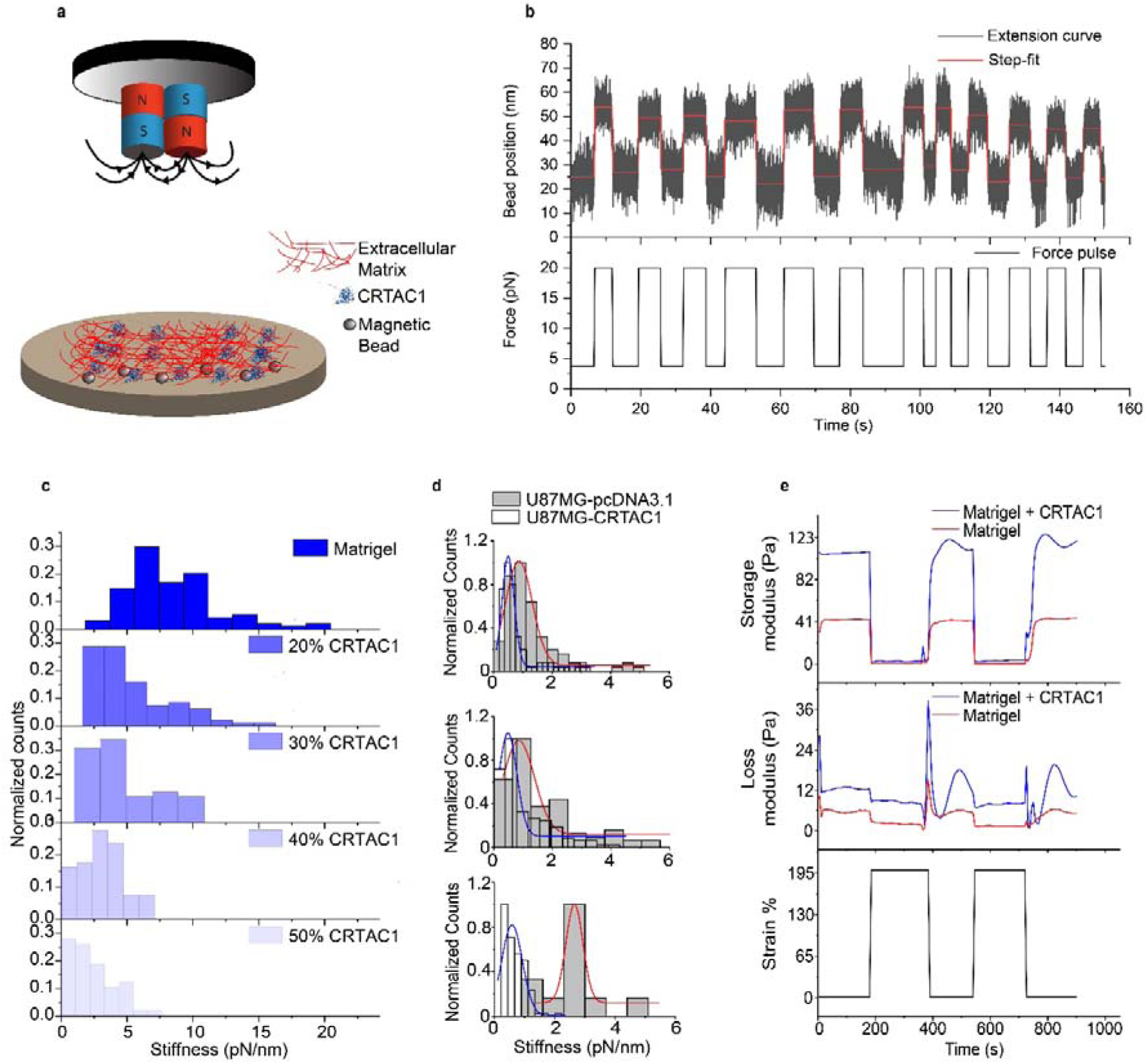
Stiffness and elasticity changes by CRTAC1 in the ECM. (**a**) Schematic of experiment on our lab made magnetic tweezers. **(b)** A representative extension of bead position and force pulse. The red line maps the steps obtained from Autostepfinder. (**c**) Different concentrations (0, 20%, 30%, 40% and 50%) of CRTAC1 were used compared to Matrigel and Collagen I thereby decreasing stiffness as concentration of CRTAC1 was increasing. **(d)** The extension of magnetic beads in Z direction was measured, and stiffness was calculated at 24, 48 and 72 hours after transfecting U87MG cells with CRTAC1 and empty vector as a control. (**e**) Elasticity measurements were done by using rheology. 200% strain was applied and released every 200s upto 1000 s. Elasticity was higher by approx. 70 Pa in presence of CRTAC1 suggesting crosslinked ECM in its presence

Incorporation of CRTAC1-A reduced bulk ECM stiffness nearly two-fold compared with control matrices. This reduction was concentration dependent, with progressive decreases observed as CRTAC1-A supplementation increased from 0% to 50% (Fig. 6c). To assess whether similar effects occur in cell-derived matrices, we measured the stiffness of ECM produced by U87MG cells following CRTAC1-A overexpression. Compared with matrices generated by control cells, CRTAC1-A–expressing cells produced ECM with ∼75% lower stiffness over time (Fig. 6d). Independent validation using parallel-plate rheometry confirmed reduced stiffness in CRTAC1-A–supplemented matrices (Supplementary Fig. 5a,b), and similar concentration-dependent softening was observed in both polyacrylamide and collagen gels (Supplementary Fig. 6a,b).

Because bulk stiffness does not fully capture nonlinear mechanical behavior under large deformations, we next examined viscoelastic recovery using time-resolved oscillatory rheology. Matrices were subjected to cyclic strain protocols alternating between low strain (within the linear viscoelastic regime) and high strain (∼200%). Under low-strain conditions, CRTAC1-A–supplemented Matrigel exhibited a modestly higher storage modulus (G′) compared with Matrigel alone (Fig. 6e), indicating enhanced network integrity under small deformations. Transition to high strain resulted in marked decreases in both storage (G′) and loss (G″) moduli in both conditions, consistent with network yielding.

Strikingly, upon returning to low strain, CRTAC1-A–containing matrices demonstrated rapid and reproducible recovery of G′ across successive strain cycles, whereas control matrices recovered to a lower plateau (Fig. 6e; Supplementary Fig. 7). The loss modulus (G″) in CRTAC1-A–supplemented matrices showed transient increases during high-strain intervals, indicative of enhanced reversible energy dissipation.

Together, these data indicate that CRTAC1-A reduces bulk stiffness under static loading while enhancing nonlinear elastic recovery and reversible dissipation under cyclic deformation. Thus, CRTAC1-A remodels collagen-rich ECM into a mechanically compliant yet structurally resilient network that resists irreversible plastic remodeling rather than simply stiffening or softening the matrix.

### CRTAC1-A–mediated ECM remodeling enhances temozolomide sensitivity

Given that CRTAC1-A remodels collagen-rich matrices into a mechanically compliant yet resilient state, we next investigated whether this altered ECM context influences responsiveness to temozolomide (TMZ), the standard chemotherapeutic agent used in GBM treatment.

U87MG spheroids were embedded in collagen I or Matrigel matrices in the presence or absence of recombinant CRTAC1-A. CRTAC1-A was supplemented at a 1:1 molar ratio relative to collagen I unless otherwise specified. Control matrices received vehicle (HEPES buffer) alone. Spheroids were treated with 1 mM TMZ, and cell death was assessed in a time-dependent manner (Fig. 7a). CRTAC1-A–supplemented spheroids exhibited significantly increased cell death compared with control spheroids at 24 h (+9.8%, p = 0.0002), 48 h (+8.8%, p = 0.0021), and 72 h (+13.2%, p < 0.0001) following TMZ treatment (Fig. 7b).

**Fig 7:**
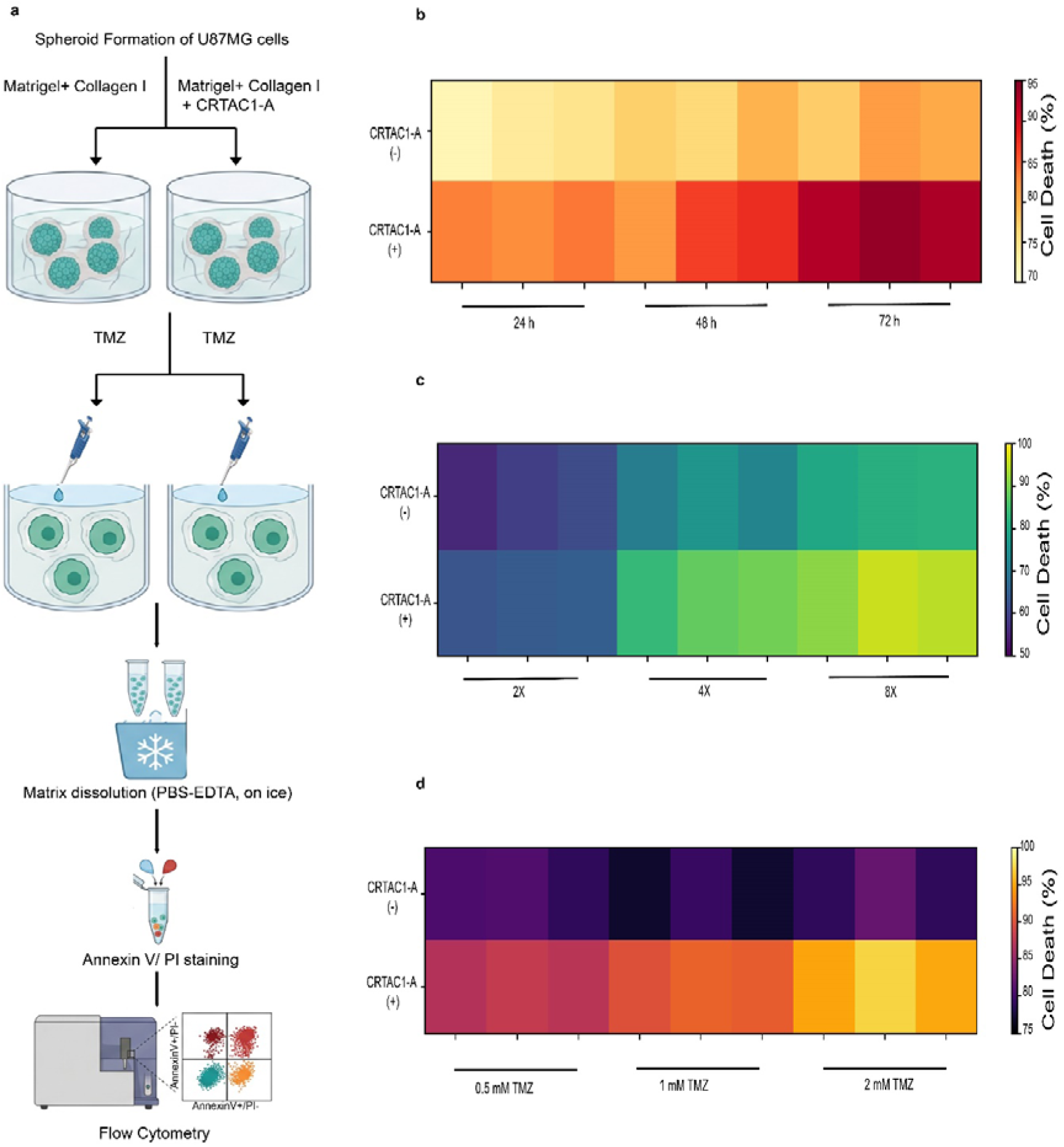
CRTAC1-A sensitizes U87MG spheroids to Temozolomide (TMZ) induced cytotoxicity. (**a)** Schematic illustration of the 3D experimental model using U87MG spheroids embedded in a composite Matrigel and Collagen I ECM supplemented with recombinant CRTAC1-A. (**b)** Time-course analysis of cell death in U87MG spheroids (n = 4) within a CRTAC1-A enriched ECM. Significant potentiation of apoptosis was observed at 24, 48, and 72 hours post-treatment (p = 0.0002, 0.0021 and < 0.0001, respectively). (**c)** Dose-dependent effect of CRTAC1-A concentration on TMZ mediated cytotoxicity. Spheroids treated with 1 mM TMZ showed significantly increased cell death in the presence of higher CRTAC1-A titers (2x, 4x, and 8x concentrations; p=0.0360, < 0.0001, and < 0.0001, respectively). (**d)** Optimization of TMZ dosage (0.5, 1, and 2 mM) in U87MG spheroids (n = 4), identifying 1 mM as the optimal concentration for subsequent synergistic studies (p = 0.0004, 0.0001, and 0.0003, respectively). Quantitative cell death was determined via flow cytometry. Statistical significance was assessed using a two-tailed Mann-Whitney test.

To determine whether this sensitization effect is concentration dependent, Matrigel-embedded spheroids were supplemented with increasing CRTAC1-A concentrations (2x, 4x, and 8x molar ratio relative to collagen I). Enhanced TMZ-induced cell death was observed at 2x (+5.3%, p = 0.0360), 4x (+13.7%, p < 0.0001), and 8x (+13.0%, p < 0.0001) CRTAC1-A supplementation compared with controls (Fig. 7c).

We further evaluated whether CRTAC1-A enhances responsiveness across increasing TMZ doses. At 0.5 mM TMZ, CRTAC1-A supplementation increased cell death by 7.8% (p = 0.0004); at 1 mM TMZ by 13.2% (p = 0.0001); and at 2 mM TMZ by 17% (p = 0.0003) relative to control matrices (Fig. 7d).

Together, these results indicate that CRTAC1-A–mediated ECM remodeling enhances TMZ-induced cytotoxicity in a time- and dose-dependent manner. These findings are consistent with the notion that viscoelastic normalization of the ECM may reduce protective invasive remodeling and improve therapeutic responsiveness.

## Discussion

GBM remains one of the most lethal malignancies, driven by diffuse invasion and a highly adaptive tumor microenvironment. Increasing evidence establishes the ECM as an active regulator of tumor progression rather than a passive scaffold, integrating biochemical signalling with mechanical cues that govern proliferation, invasion, and therapeutic response. In this study, we identify CRTAC1-A as an ECM-associated regulator that modulates collagen architecture and viscoelastic behavior in a manner that constrains glioma aggressiveness.

ECM stiffening is widely recognized as a hallmark of GBM progression and has been linked to enhanced integrin clustering^20^, cytoskeletal contractility^21^, and mechanotransductive^22^ signaling that enable persistent matrix remodeling and invasion^23^. Our findings refine this paradigm by demonstrating that malignant progression cannot be explained by stiffness alone. Instead, CRTAC1-A reprograms matrix mechanics toward a compliant yet plasticity-resistant state. Through direct interaction with collagen I, CRTAC1-A promotes fibril bundling and junctional connectivity while preserving fibril ultrastructure. This structural reorganization reduces bulk stiffness under static loading yet enhances nonlinear elastic recovery and reversible energy dissipation under cyclic strain. Importantly, such matrices resist irreversible deformation despite appearing softer under small perturbations. These findings indicate that invasion depends not only on absolute stiffness but critically on matrix plasticity, the capacity for permanent remodeling under cellular forces^24,34^.

Clinically, *CRTAC1* expression is elevated in LGGs and declines progressively with tumor grade. Isoform-specific analyses reveal that this enrichment is driven predominantly by the extracellular CRTAC1-A isoform, whereas the brain-enriched CRTAC1-B isoform is reduced. Functional assays demonstrate that CRTAC1-A restrains glioma proliferation and invasion in three-dimensional matrices, consistent with its association with improved patient survival. Similar tumor-restraining roles for *CRTAC1* have been reported in lung and urothelial cancers^25,26^, suggesting a broader role for this protein family in regulating tumor–ECM interactions.

Alternative splicing emerges as a key regulatory mechanism shaping CRTAC1-A isoform composition in gliomas. We demonstrate that exon inclusion favoring CRTAC1-A is enhanced in tumor tissues and increases with grade, indicating dynamic isoform regulation during disease progression. Notably, this splicing shift occurs independently of IDH mutational status, suggesting regulation by splicing-factor–mediated mechanisms rather than direct IDH-driven epigenetic control. Given the widespread dysregulation of splicing programs in gliomas^27^, altered isoform balance may represent an underappreciated determinant of tumor–matrix signalling.

While CRTAC1-A exon inclusion increases with tumor grade, total transcript abundance declines in GBM. Our data indicate that this inverse expression pattern arises from stage-dependent genomic and epigenetic mechanisms. IDH1-mutant LGGs exhibit reduced CRTAC1-A promoter methylation and elevated transcript levels, consistent with locus-specific redistribution of methylation marks reported in IDH1-mutant tumors^29,30,31^. In contrast, progressive shallow deletions of the *CRTAC1* locus occur in GBM and coincide with reduced expression. Because chromosome 10q loss of heterozygosity is a frequent event in GBM^32,33^ and *CRTAC1* resides at 10q24.2, monoallelic deletion likely contributes to reduced *CRTAC1* expression in advanced disease. Thus, distinct regulatory layers including alternative splicing, epigenetic modulation, and copy-number alteration collectively shape CRTAC1-A isoform abundance across glioma grades.

Mechanistically, reduced *CRTAC1* expression in GBM correlates with increased ECM rigidity and enhanced invasive potential. Notably, CRTAC1-A mediated remodeling did not alter phosphorylation of the mechanotransduction marker FAK (Supplementary Fig. 8), suggesting that its invasion-suppressive effects arise primarily from modulation of matrix plasticity rather than direct suppression of canonical integrin signalling. By reorganizing collagen networks through reversible interactions rather than permanent crosslinking, CRTAC1-A generates ECMs that dissipate mechanical energy and recover rapidly after deformation. Such environments may limit the stabilization of invasion tracks required for persistent collective migration.

CRTAC1-A therefore represents a distinct class of matricellular regulator that modulates ECM mechanics without functioning as a primary load-bearing component. Similar to netrin-4, which reduces tumor invasiveness by altering laminin network organization^34^, CRTAC1-A reprograms collagen architecture to constrain malignant behavior. In contrast to stiffness-promoting ECM proteins such as tenascin-C, whose elevated expression correlates with poor prognosis in gliomas^25^, CRTAC1-A underscores that tumor aggressiveness is governed not simply by matrix rigidity but by dynamic regulation of force transmission and irreversible remodeling.

An important consideration is that CRTAC1-A appears to be a human-specific isoform and is not conserved in commonly used rodent models such as mouse or rat. This species specificity limits the feasibility of conventional murine glioma models for in vivo validation. While future studies employing humanized systems or patient-derived xenograft models will be necessary to assess physiological relevance in vivo, the absence of a conserved ortholog underscores the possibility that CRTAC1-A represents an evolutionarily specialized regulator of human ECM architecture. This highlights a broader limitation of traditional rodent systems in modeling certain aspects of human tumor–matrix regulation.

Beyond its mechanistic role, CRTAC1-A may also hold clinical relevance. Its isoform-specific enrichment in LGG, association with IDH1 mutation status, and correlation with patient survival suggest potential utility in prognostic stratification. However, further clinical validation will be necessary to determine whether CRTAC1-A expression can serve as a predictive biomarker or therapeutic target.

In summary, our findings identify CRTAC1-A as a reversible matricellular regulator that reprograms tumor ECM viscoelasticity. By shifting collagen networks toward a compliant yet plasticity-resistant state, CRTAC1-A constrains glioma invasion and enhances therapeutic responsiveness. These results expand the stiffness-centric view of tumor mechanobiology and establish dynamic mechanical regulation as a critical determinant of glioma progression.

## Conclusion

ECM-mediated resistance remains a major obstacle in GBM therapy, as stiff and poorly adaptable matrices hinder both immune infiltration and effective drug delivery. By promoting a mechanically normalized ECM that suppresses invasion without excessive densification, CRTAC1-A represents a promising target for matrix-directed therapies. Modulating CRTAC1-A activity in combination with standard treatments such as temozolomide may improve therapeutic efficacy by simultaneously restraining tumor invasion and preserving ECM permeability. Future studies using preclinical GBM models will be crucial to assess whether targeting CRTAC1-A can reprogram the tumor microenvironment toward a less aggressive, more therapy-responsive state. Collectively, our work highlights the therapeutic potential of reversible ECM regulators and advances a mechanobiology-driven framework for GBM treatment.

## Materials and Methods

### Cell Culture and Transfection

U87MG and HEK293T cell lines were obtained from NCCS, Pune and were cultured in Dulbecco’s Modified Eagle’s Medium (AL068A-500ML, Himedia) (DMEM) supplemented with 5% fetal bovine serum (FBS) (RM9955-500ML, Himedia) 100 U/mL penicillin, and 100 μg/mL streptomycin at 37°C in a humidified atmosphere containing 5% CO_2_. Transfection of the plasmid DNA into HEK293T cells with Ex-Cell media (14366C-1000ML, Sigma) was achieved using polyethyleneimine (Sigma) according to the manufacturer’s protocol.

### Cloning of CRTAC1-A and stable CRTAC1-A U87MG generation

Human CRTAC1-A coding sequence was cloned into a pcDNA-3.1 vector. CRTAC1-A in pDONR221 (plasmid id - HsCD00744950) vector was purchased from DNASU plasmid repository. The sequence was subcloned into a pcDNA-3.1 vector using primers (Supplementary Table 2) carrying Nde1 and BamH1 cloning sites, with His tag sequence added in-frame on the C-terminus after the coding sequence to generate pcDNA-3.1-CRTAC1-His. Cells were seeded in tissue culture-treated flasks and grown to approximately 70-80% confluency prior to transfection. U87MG-CRTAC1 and control stable cells were generated by transfection of pcDNA3.1-CRTAC1-His construct and empty vector pcDNA3.1, followed by selection in media supplemented with 100 µg/ml G418 antibiotic. pcDNA3.1-CRTAC1-SGH clone was generated by cloning CRTAC1-A sequence into pcDNA3.1-GFP vector for fluorescence studies. (Supplementary Table 2)

### Protein Expression and Purification

Ex-Cell media from transfected cells was harvested at 48 h post-transfection and dialyzed against lysis buffer (50 mM Tris-HCl pH 8.0, 150 mM NaCl, 1 mM EDTA, 1% Triton X-100). After dialysis it was applied to a pre-equilibrated Ni-NTA affinity chromatography column (HisTrap HP, GE Healthcare) equilibrated with the lysis buffer. Unbound proteins were removed by washing with the wash buffer (50 mM Tris-HCl pH 8.0, 300 mM NaCl, 20 mM imidazole). The target protein was eluted with the elution buffer (50 mM Tris-HCl pH 8.0, 300 mM NaCl, 250 mM imidazole). Eluted protein fractions were collected and analysed for purity using SDS-PAGE.

### Cell proliferation Assay

Stable transfected U87MG glioblastoma cells with pcDNA3.1-CRTAC1-A and empty pcDNA3.1 were cultured in DMEM supplemented with 10% FBS and 1% penicillin-streptomycin under standard conditions (37°C, 5% CO2). Cells were seeded in 96-well plates at a density of 3000 cells per well and allowed to adhere overnight. Incubation was done for 24,48,72,96 and 120 h to allow cell proliferation. Following this, 20 µL of MTT solution (5 mg/mL in PBS) was added to each well, and cells were incubated at 37°C for 2 h allowing metabolically active cells to reduce the yellow MTT tetrazolium salt to purple formazan crystals. After incubation, the medium was carefully removed, and 100 µL of dimethyl sulfoxide (DMSO) was added to each well to dissolve the formazan crystals. The absorbance was measured at 570 nm using a microplate reader to quantify cell viability and proliferation. The proliferation rate of the CRTAC1-A transfected U87MG cells was calculated relative to control cells.

### Spheroid Invasion Assay

U87MG glioblastoma spheroids were generated using the hanging drop method. Briefly, 20□µl droplets of DMEM supplemented with 5% FBS, each containing 5000 cells, were deposited onto the inner lids of 10□cm culture dishes. To prevent evaporation, lids were inverted over dishes containing 1× PBS and incubated for 120□h to allow for spontaneous spherical aggregation. Following maturation, individual spheroids were harvested and embedded in a 20□µl ECM scaffold. This scaffold consisted of either a control matrix—comprising Matrigel (4□mg/ml; Corning, 354234) and collagen I (0.5□mg/ml; Sigma-Aldrich, C3867) or an experimental ^matrix^ further supplemented with 70□µM CRTAC1-A. To ensure centralized positioning of the spheroids within the 3D volume, plates were incubated in an inverted position at 37□°C for 1□h to facilitate matrix polymerization. Upon solidification, the matrices were overlaid with DMEM containing 5% FBS. Spheroid invasion into the surrounding ECM was monitored via bright-field microscopy using a Leica DMi8 platform at 4× magnification. Images were captured at 24□h intervals over a 120 h period. The total area of invasion was quantified using ImageJ software (version 2.1.0/1.53c), with the primary spheroid body serving as the baseline for radial expansion.

### Pull down Assay

To investigate the interaction between CRTAC1-A and Collagen I, a co-immunoprecipitation pull-down assay was performed using immobilized antibodies. Protein A/G beads were first equilibrated by washing three times in ice-cold PBS supplemented with a protease inhibitor cocktail. For antibody immobilization, 2□µg of purified anti-CRTAC1-A antibody was conjugated to the beads in 500□µl of binding buffer (20□mM Tris-HCl, pH 7.4; 150□mM NaCl; 0.1% Tween-20) for 2□h at 4□°C under constant, gentle rotation. The antibody-bead complexes were then incubated with purified CRTAC1-A protein for 4□h at 4□°C. To assess binding affinity, human ECM (Sigma E0282) was added to the mixture and incubated overnight at 4□°C. Following incubation, the beads were subjected to five stringent washes with ice-cold wash buffer to remove non-specifically bound proteins. The target protein complexes were eluted using 50□µl of elution buffer (0.1□M glycine, pH 2.5). To maintain protein stability and prevent acid-mediated degradation, the eluates were immediately neutralized with 1□M Tris-HCl (pH 8.0) followed by immunoblot with CRTAC1-A and Collagen I antibody.

### Co-localization Assay

For colocalization analysis, 30□mm glass-bottom dishes were functionalized with Cy5-labeled Collagen I. U87MG cells were seeded onto these substrates and subsequently transfected with plasmids encoding GFP-tagged CRTAC1-A, following the manufacturer’s optimized protocol. 48 h post-transfection, the culture medium was aspirated, and the cells were washed with PBS before being fixed in 4% (w/v) paraformaldehyde (PFA) for 15□min at room temperature. High-resolution fluorescence imaging was performed using a Zeiss LSM 980 confocal laser scanning microscope (CLSM), equipped with appropriate laser lines for GFP and Cy5 excitation.

### Gene Expression data

mRNA expression of overexpressed and downregulated genes and overall patient survival data in LGG and GBM patients was extracted from GEPIA2 webserver^29^. GEPIA2 surveys the gene expression data from TCGA (The Cancer Genome Atlas) and GTEx (Genotype-Tissue Expression) portals^35^. The genes with significant differential expression (p-value<0.05) among LGG and GBM were identified. DNA methylation and copy number variations data concerning CRTAC1 expression was obtained from the cBioportal database. To validate CRTAC1 expression, U87MG cells were transfected with IDH1-R132H mutant (Addgene id-62907) and IDH1-WT (Addgene id-62906) constructs. pcDNA3-Flag-IDH1-R132H (Addgene id-62907) and pcDNA3-Flag-IDH1 (Addgene id-62906) were gifted by Yue Xiong^36^. After 48 hours cDNA was synthesised from RNA extracted from transfected cells and real time PCR was carried out using CRTAC1 specific primers and 18S rRNA specific primers for reference control (Supplementary Table 3).

### Alternative Splicing Data Acquisition and Processing

To investigate the alternative splicing landscape of CRTAC1-A, we utilized two comprehensive cancer splicing resources: TCGASpliceSeq and OncoSplicing. SpliceSeq data, derived from The Cancer Genome Atlas (TCGA) RNA-seq cohorts, were used to extract Percent Spliced In (PSI) values, which represent the ratio of transcript isoforms including a specific exon over the total number of transcripts. Splicing profiles were cross-referenced with the OncoSplicing database to ensure robust quantification of isoform variations across glioblastoma multiforme (GBM) and lower-grade glioma (LGG) samples. The distribution of PSI values across the normal-LGG-GBM progression was visualized using box-and-whisker plots.

### Rheological Measurements

The viscoelastic properties of the Matrigel–Collagen I matrices were characterized using a Anton Paar Rheometer 302 equipped with a 25 mm diameter parallel-plate geometry. To ensure uniform matrix assembly, 500□µl of the ECM mixture comprising Matrigel (4.5□mg/ml) and Collagen I (0.5□mg/ml), with or without the addition of CRTAC1-A was loaded onto the lower plate pre-cooled to 4□°C. The measurement gap was set to 0.5□mm. To define the linear viscoelastic region (LVR), an amplitude sweep was first conducted by varying the oscillatory strain from 0.01% to 100% at a constant frequency of 1□Hz. Subsequent frequency sweeps were performed between 1 and 30□Hz at a fixed strain of 0.1% (within the LVR) at 37□°C for 30□min to monitor the polymerization kinetics and steady-state moduli^37^. The structural robustness and recovery potential of the hydrogels were evaluated through step-strain experiments. The storage modulus (G’) and loss modulus (G’’) were recorded over a 1000□s period by alternating between low-strain (1%) and high-strain (200%) conditions every 200□s. This cyclic deformation protocol was used to quantify the impact of CRTAC1-A on the mechanical integrity and recovery of the ECM.

### Stiffness measurement by Magnetic Tweezers

To characterize the micromechanical properties of the extracellular matrix (ECM) and the forces exerted by U87MG cells, glass coverslips were chemically activated through immersion in piranha solution for 2□h. Following treatment, coverslips were rinsed in distilled water and subjected to five cycles of bath sonication to ensure a contaminant-free surface. For force probe immobilization, 100□µl of magnetic beads were deposited onto the air-dried coverslips and incubated at room temperature for 1□h. A Collagen I scaffold (3□mg/ml) was prepared by 60-fold dilution in 20□mM acetic acid and allowed to polymerize over the bead-coated surface for 1□h at 37□°C. U87MG cells (1 x 10^5^ cells/coverslip) were then seeded onto the collagen matrix. Transient transfection with the pcDNA3.1-CRTAC1 plasmid was performed at 24□h intervals over a total period of 72□h to maintain stable expression levels. For *in vitro* protein-level studies, recombinantly purified CRTAC1-A was incorporated into a Matrigel–Collagen I matrix at a 2:1 ratio (v/v). This composite gel was allowed to polymerize on bead-coated coverslips at 37□°C for 1□h. Micromechanical measurements were conducted using a custom-built magnetic tweezers platform. Mechanical stiffness and extension profiles were recorded by applying a constant force of 100□pN for cellular measurements and 20□pN for purified protein-matrix studies. To determine the probability distribution of the resulting displacement data, trajectories were analyzed using kernel density estimation^38^. All data fitting and visualization were performed using OriginPro software.

### Transmission Electron Microscopy

Collagen I solutions (1 mg/mL in 0.1 M acetic acid) were dialyzed against 0.1 M PBS (pH 7.4) at 4°C for 24 h to neutralize and promote fibrillogenesis in presence or absence of CRTAC1-A. Aliquots (5 µL) were applied to glow-discharged Formvar-carbon coated copper grids (200 mesh) for 2 min and grids were air-dried at room temperature. Imaging was performed using a JEOL JEM-F2000 TEM operated at 80 kV. Grids were loaded into a side-entry goniometer stage, and bright-field images were captured using a high-resolution Gatan One-view camera. Digital images were processed using ImageJ software for contrast enhancement and measurement of fibril alignment was done using the Orientation J plugin of ImageJ software.

### Cell death due to Temozolomide drug measured by FACS

Spheroids were established and embedded in a matrix of 4 mg/mL Matrigel and 0.5 mg/mL Collagen I. To evaluate the potential of CRTAC1-A to enhance temozolomide (TMZ) efficacy, embedded spheroids were cultured in DMEM supplemented with TMZ. Following incubation, the matrix was dissociated on ice using a PBS-EDTA solution to obtain a single-cell suspension, as previously described^39^. Cell viability and apoptosis were quantified via Annexin V and Propidium Iodide (PI) staining, with data acquisition performed on a BD Aria Fusion flow cytometry.

### Preparation of stiff polyacrylamide gels

Thin polyacrylamide (PAA) gels with a target stiffness of 5 kPa were synthesized by free-radical polymerization of acrylamide and N,N’-methylenebisacrylamide (bis-acrylamide) (Sigma-Aldrich). Briefly, glass coverslips were chemically functionalized to ensure covalent attachment of the gel. 3-aminopropyltriethoxysilane (APTES) was applied to the bottom coverslips for 5 minutes, followed by treatment with 0.5% glutaraldehyde for 30 minutes.To achieve a stiffness of 5 kPa, a precursor solution was prepared containing 5% (w/v) acrylamide and 0.15% (w/v) bis-acrylamide in deionized water. The mixture was degassed for 15 minutes to prevent oxygen inhibition of the polymerization process. Polymerization was initiated by the addition of 10% (w/v) ammonium persulfate (APS) at a 1:100 v/v ratio and N,N,N’,N’-tetramethylethylenediamine (TEMED) at a 1:1000 v/v ratio^40^. A volume of 25 μl of the mixture was sandwiched between the functionalized coverslip and a top coverslip to ensure hydrophobicity. Gels were allowed to polymerize for 45 minutes at room temperature in a humidified chamber. After polymerization, the top coverslip was carefully removed, and the gels were washed extensively in phosphate-buffered saline (PBS) to remove unreacted monomers. To facilitate cell adhesion, PAA gels were functionalized using the linker NHS-AA ester (Thermo Fisher Scientific). A 0.5 mg/ml solution in HEPES buffer (50 mM, pH 8.5) was applied to the gel surface. Following two washes with HEPES, the gels were incubated with Type I Collagen (control) and with equal amounts of CRTAC1-A overnight at 4□ to provide an extracellular matrix (ECM) interface for cell seeding^41^.

### Statistical Analysis

All data are expressed as mean ± standard deviation (SD) or median depending on data distribution. Comparisons between two groups were performed using the unpaired two-tailed Student’s t-test or Mann-Whitney U test for parametric and non-parametric data, respectively. For multiple group comparisons, one-way analysis of variance (ANOVA) with appropriate post hoc tests or Kruskal-Wallis test was used. Correlation analyses were performed using Pearson’s correlation coefficients as appropriate. Statistical significance was set at a two-sided p value < 0.05. Statistical analyses were conducted using GraphPad Prism software version 8.

## Acknowledgements

We acknowledge Department of Biotechnology, Government of India for funding the development of Magnetic Tweezers (BT/PR55022/BMS/85/563/2024). We also thank IISER Mohali for the Institute Notional grant. We acknowledge the Department of Biological Science (DBS), IISER Mohali for the Super resolution microscopy facility supported by DST-FIST grant (SR/FST/LS-II/2017/097). We also acknowledge the DBS, IISER Mohali for flow cytometer facility. V.S. acknowledges IISER Mohali for Postdoctoral research fellowship. D.S., P.S. and D.N. acknowledge CSIR for fellowship. We thankfully acknowledge DNASU plasmid repository, Arizona USA for providing CRTAC1 plasmid (plasmid id - HsCD00744950). We acknowledge the use of Perplexity AI during editing to improve the clarity and grammatical accuracy of the text.

## Author information

### Contribution

S.R. conceptualized the work. V.S. designed, performed, and analyzed all the cellular experiments as well as TCGA data analysis including IDH1 regulation of CRTAC1. V.S. performed and analyzed flow cytometry and qPCR data. D.S. cloned and purified the protein. D.S. performed and analyzed all the imaging experiments as well as pull-down data. D.S. prepared samples for stiffness experiments. P.S. performed and analyzed magnetic tweezers data. D.N. performed and analyzed rheological measurements. N.D. performed differential gene expression between LGG and GBM. S.R., V.S., and D.S. drafted the manuscript. S.R, V.S., D.S and P.S edited the manuscript.

## Declarations

Authors declare no competing interests

## Data Availability Statement

Data will be available from the corresponding author upon reasonable request.

