## Supplementary Information for "CRTAC1-A reprograms extracellular matrix viscoelasticity to constrain glioma progression"

List of Supplementary Figures and Tables

| **Fig 1** | Integrative bioinformatics pipeline for the identification and functional prioritization of *CRTAC1.* |
| --- | --- |
| **Fig 2** | CRTAC1-A transcript abundance is independent of IDH mutational status in LGG. |
| **Fig 3** | *In vitro* pull-down assay validating the interaction between CRTAC1-A and Collagen I. |
| **Fig 4** | Mechanical characterization of ECM stiffness via magnetic tweezers and automated bead tracking. |
| **Fig 5** | Rheological characterization of CRTAC1-A-mediated modulation of Matrigel mechanics. |
| **Fig 6** | CRTAC1-A mediated attenuation of hybrid and collagen-only matrix mechanics. |
| **Fig 7** | Cyclic strain-recovery analysis of CRTAC1-A supplemented composite matrices. |
| **Table 1** | List of genes differentially expressed between LGG and GBM. |
| **Table 2** | List of primers for cloning. |
| **Table 3** | List of primers for qPCR. |

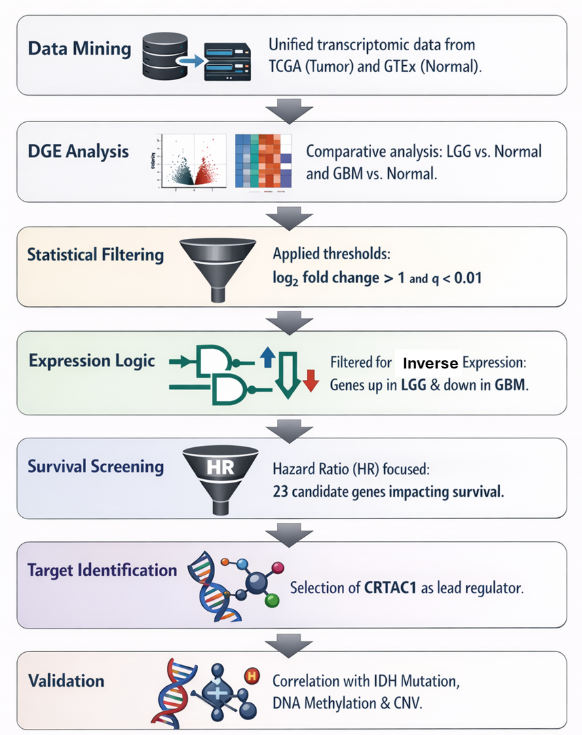

**Supplementary Fig 1. Integrative bioinformatics pipeline for the identification and functional prioritization of *CRTAC1*.** Schematic overview of the *in silico* strategy employed to identify *CRTAC1* as a candidate gene for experimental work in glioma.

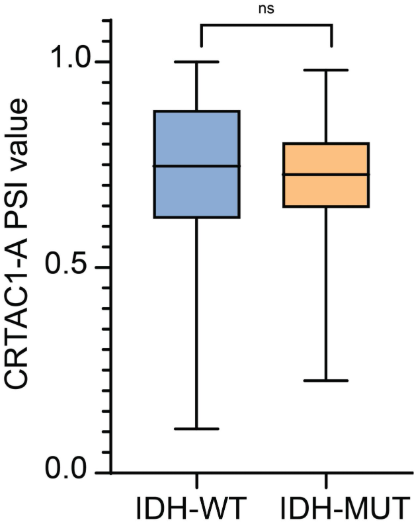

**Supplementary Fig 2. CRTAC1-A transcript abundance is independent of IDH mutational status in LGG.** Analysis of Percent Spliced-In (PSI) values for the CRTAC1-A isoform, stratified by IDH genotype in the LGG cohort. The data demonstrate that CRTAC1-A enrichment remains invariant across IDH mutational subtypes, suggesting that the observed isoform switching is governed by mechanisms distinct from IDH-mediated epigenetic remodeling. Statistical significance was assessed using a two-tailed Mann-Whitney test (p > 0.05).

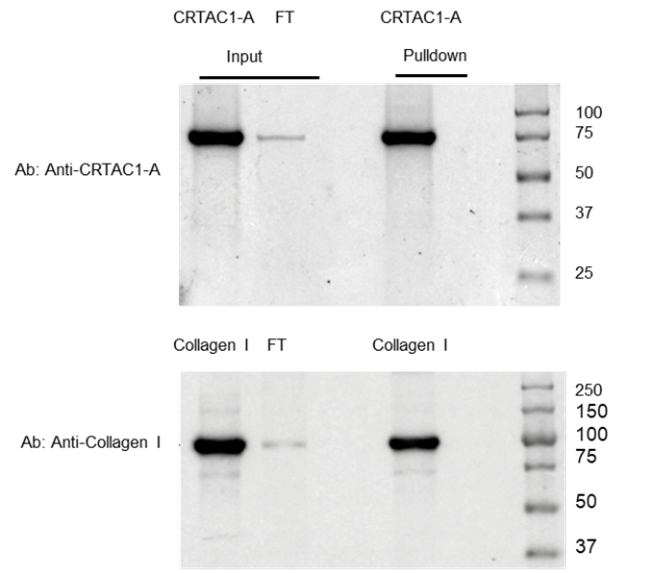

**Supplementary Fig 3. *In vitro* pull-down assay validating the interaction between CRTAC1-A and Collagen I.** Co-immunoprecipitation was performed using purified CRTAC1-A incubated with a Matrigel+Collagen I composite matrix for 1 h at room temperature. The complex was captured using an anti-CRTAC1-A antibody immobilized on Protein A/G agarose beads through an overnight incubation at 4 °C. Representative western blots of the eluted fractions demonstrate the co-enrichment of Collagen I alongside the CRTAC1-A bait, confirming physical association between the two proteins.

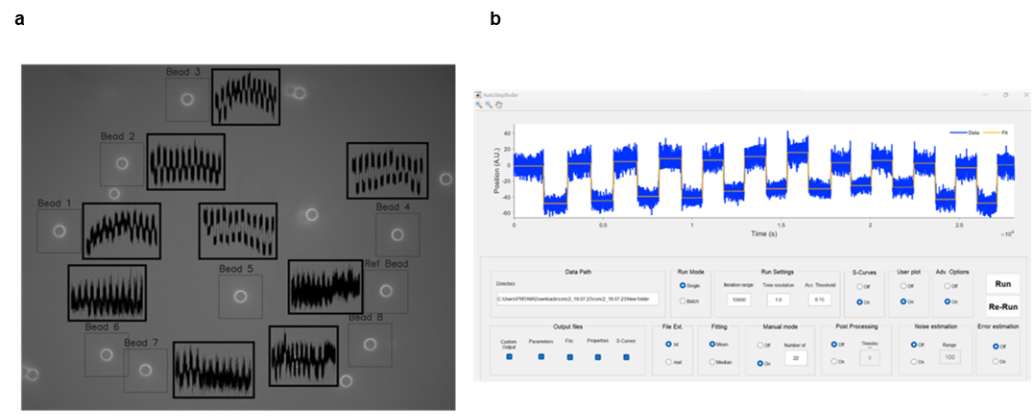

**Supplementary Fig 4. Mechanical characterization of ECM stiffness via magnetic tweezers and automated bead tracking. (a)** Representative micrograph (100× magnification) of paramagnetic beads embedded within a composite Matrigel+Collagen I matrix. Local mechanical properties were assessed using a custom Python based multi-bead tracking algorithm; overlaid displacement trajectories illustrate the elastic response of the matrix to oscillatory magnetic force. The displacement amplitude is inversely proportional to stiffness. (**b)** Implementation of the Autostepfinder (MATLAB) software for the objective detection of discrete positional transitions. The platform utilizes a least-mean-squares optimization algorithm to perform automated step-fitting on time-extension trajectories derived from force-clamp or oscillatory traces, enabling precise quantification of stochastic displacement events within ECM.

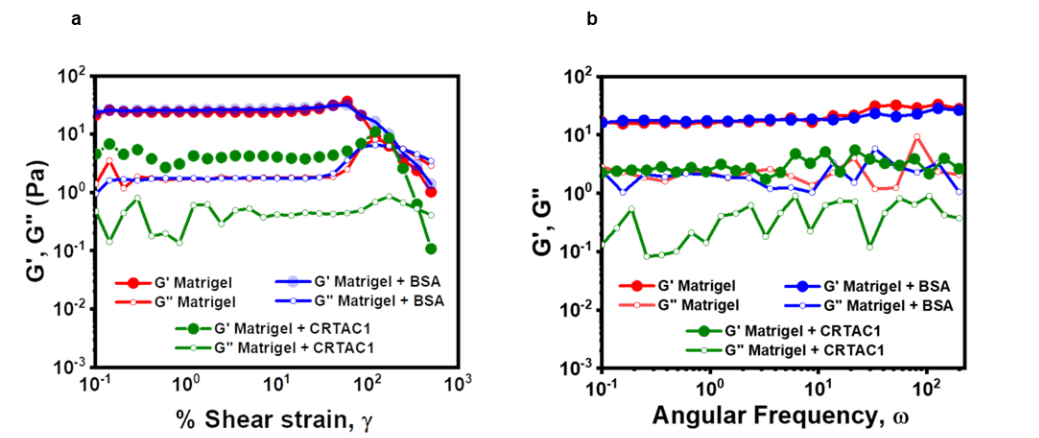

**Supplementary Fig 5. Rheological characterization of CRTAC1-A-mediated modulation of Matrigel mechanics.** (**a)** Strain-amplitude sweep analysis of Matrigel under control (untreated and BSA-supplemented) and CRTAC1-A treated conditions. The storage modulus (G') and loss modulus (G'') are plotted as a function of oscillatory strain amplitude. Control samples exhibit a robust linear viscoelastic region with a higher G', indicative of superior mechanical rigidity. Incorporation of CRTAC1-A results in a significant reduction in G' across the strain range, signifying a weakened elastic network. The G'/G'' crossover point represents the yield strain, marking the transition from an elastic solid-like to a viscous liquid-like state. (**b)** Frequency sweep profiles (0.1–10 rad/s) illustrating the dynamic mechanical response of the matrix. Control and BSA-treated Matrigel maintain consistently higher G' values compared to CRTAC1-A supplemented gels, demonstrating that the reduction in stiffness is independent of the deformation timescale.

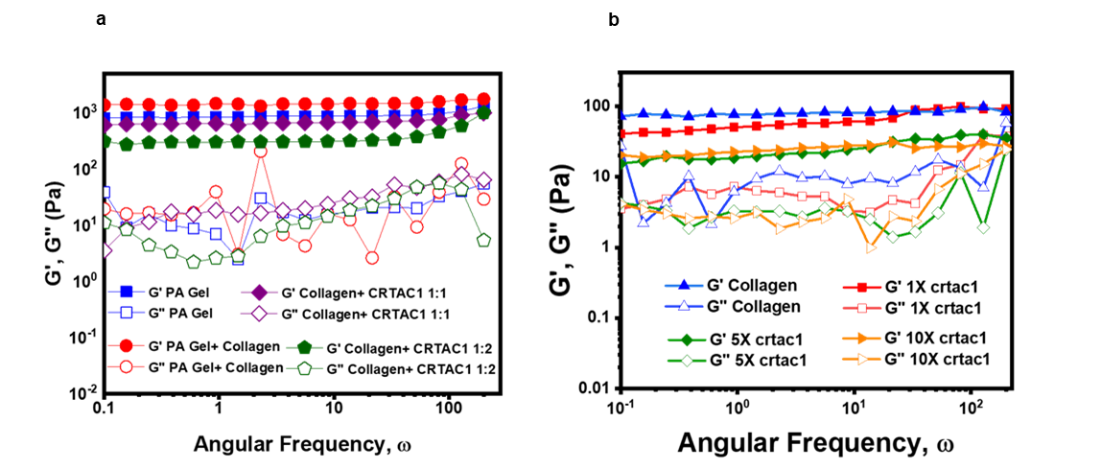

**Supplementary Fig 6. CRTAC1-A mediated attenuation of hybrid and collagen-only matrix mechanics. (a)** Rheological characterization of polyacrylamide (PA) gels functionalized with Collagen I. Storage (G') and loss (G'') moduli were determined via frequency sweep measurements at 37 °C (0.01 strain amplitude, parallel plate geometry). While the integration of Collagen I into the PA matrix provides elastic reinforcement, the addition of CRTAC1-A prior to gelation results in a significant reduction of G', suggesting that CRTAC1-A interferes with the structural assembly of the collagen-PA network. (**b)** Dose-dependent mechanical profiles of collagen hydrogels supplemented with increasing concentrations of CRTAC1-A. Frequency sweep analysis demonstrates a progressive, concentration-dependent decrease in G' across the measured spectral range (0.1–10 rad/s). Throughout the attenuation, G' remains consistently higher than G'', confirming that the hydrogels retain their characteristic solid-like viscoelastic behavior despite reduced stiffness. These data highlight the capacity of CRTAC1-A to modulate collagen-based matrix mechanics in a dose-responsive manner.

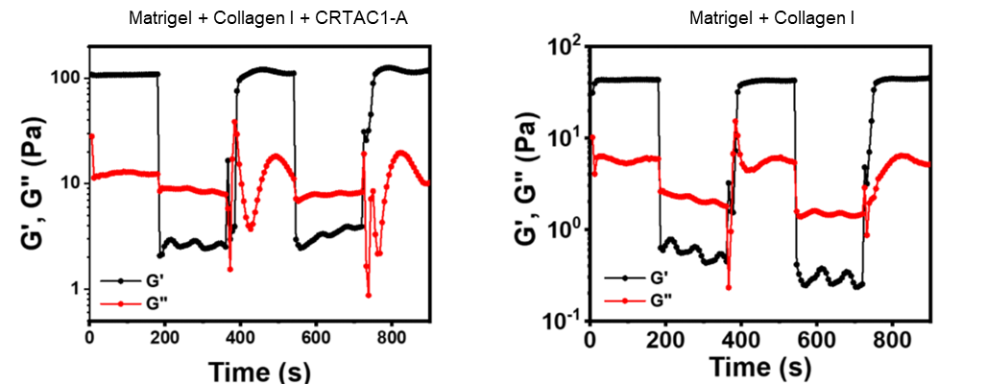

**Supplementary Fig 7. Cyclic strain-recovery analysis of CRTAC1-A supplemented composite matrices.** Dynamic mechanical response of Matrigel+Collagen I composites with and without CRTAC1-A treatment under high strain conditions. The storage (G') and loss (G'') moduli were monitored over a 1,000 s duration using a periodic step-strain protocol, alternating between a high strain state (200% for 200 s) and a recovery state (1% strain). Incorporation of CRTAC1-A into the Matrigel+Collagen I gel resulted in a marked elevation of G' during recovery phases, indicating that the protein enhances the elastic resilience and structural restoration of the composite network following large-amplitude deformation.

| **Gene Name** | **log2 fold change (LGG vs Control)** | **LGG Hazard Ratio (High)** | **p(HR)-LGG** | **log2 fold change (GBM vs Control)** | **GBM Hazard Ratio (High)** | **p(HR)-GBM** |
| --- | --- | --- | --- | --- | --- | --- |
| CHI3L1 | -1.323 | 3.3 | 3.5e−10 | 5.937 | 1.3 | 0.16 |
| COL5A2 | -1.345 | 2.5 | 1.6e−06 | 1.165 | 1.1 | 0.5 |
| FABP5 | -1.454 | 2 | 0.00012 | 2.108 | 1.6 | 0.014 |
| FABP5P7 | -1.439 | 2 | 0.00025 | 2.166 | 1.5 | 0.028 |
| G0S2 | -1.185 | 1.8 | 0.0012 | 1.668 | 1.4 | 0.069 |
| MOXD1 | -1.842 | 2 | 0.00044 | 1.628 | 1.1 | 1.1 |
| OCIAD2 | -1.622 | 2.3 | 1.1e−05 | 1.95 | 0.98 | 0.93 |
| RARRES2 | -2.267 | 2 | 0.00021 | 1.563 | 1.5 | 0.026 |
| RBP1 | -1.15 | 2.4 | 3.3e−06 | 2.668 | 1.1 | 0.55 |
| RP11-175B9.3 | -1.218 | 0.89 | 0.54 | 1.97 | 0.85 | 0.38 |
| TMBIM4 | -1.599 | 2.4 | 8.8e−06 | 3.257 | 1.3 | 0.13 |
| TMSB4XP6 | -3.254 | 5.4 | <0.0001 | 3.636 | 0.99 | 0.77 |
| CRTAC1 | 1.186 | 0.28 | 1.6e−10 | -2.006 | 1 | 0.8 |
| CSDC2 | 1.384 | 0.44 | 1.2e−05 | -1.529 | 0.96 | 0.81 |
| CSMD3 | 1.102 | 0.36 | 1.5e−07 | -1.364 | 0.85 | 0.38 |
| DSCAML1 | 1.421 | 0.43 | 5.4e−06 | -1.082 | 0.99 | 0.94 |
| PCDH15 | 2.394 | 0.4 | 1.2e−06 | -1.117 | 1 | 1 |
| RP11-138I1.4 | 1.406 | 1.3 | 0.18 | -1.161 | 1.3 | 0.19 |
| SLC25A48 | 1.108 | 0.37 | 2.4e−07 | -1.536 | 0.74 | 0.1 |
| WNT7B | 1.113 | 0.51 | 0.00032 | -1.031 | 1.3 | 0.15 |
| ZDHHC22 | 1.005 | 0.38 | 3e−07 | -1.31 | 0.88 | 0.47 |
| ZNF286A | 1.13 | 0.97 | 0.86 | -1.234 | 0.92 | 0.65 |
| ZNF488 | 1.272 | 0.61 | 0.0071 | -1.312 | 0.85 | 0.36 |

**Table 1: List of genes differentially expressed in LGG and GBM cases compared to the control group.**

| **Tag** | **Primer Name** | **Sequence (5' → 3')** |
| --- | --- | --- |
| His tag | CRTAC1Nheh1F | GCTAGCATGGAGCGCTGACC |
| His tag | CRTAC1HISBamH1R | GGATCCTCAGTGGTGGTGGTGGTGGTGGCAGCTGGGCTCGCAGCTCTC |
| GFP tag | CRTAC1-GFP-FP | GGACCCAAGCTGGGTACCATGGCTCCGAGCGCTGACC |
| GFP tag | CRTAC1-GFP-RP | CTGCCTGAAACAGGATCTAGTATGGTGAGCAAGGGAGAGGAGCGCAGCTGGGCTCGCAGCTCTC |

**Table 2: List of primers used for CRTAC1-A cloning in pcDNA3.1 vector**

| **Gene Target** | **Direction** | **Sequence (5′→3′)** |
| --- | --- | --- |
| CRTAC1 | Forward (F) | CAACCGTGATGGCAAAGTGG |
| CRTAC1 | Reverse (R) | CTCTACGGATGACGCGGAAG |
| 18S | Forward (F) | CAGTGAAACTGCGAATGGCT |
| 18S | Reverse (R) | GATAAATGCACGCATCCCCC |

**Table 3: List of primers used for qPCR**
